# Defining the Immune Responses for SARS-CoV-2-Human Macrophage Interactions

**DOI:** 10.1101/2021.07.07.449660

**Authors:** Mai M. Abdelmoaty, Pravin Yeapuri, Jatin Machhi, Katherine E. Olson, Farah Shahjin, You Zhou, Liang Jingjing, Kabita Pandey, Arpan Acharya, Siddappa N. Byrareddy, R. Lee Mosley, Howard E. Gendelman

## Abstract

Host innate immune response follows severe acute respiratory syndrome coronavirus 2 (SARS-CoV-2) infection, and it is the driver of the acute respiratory distress syndrome (ARDS) amongst other inflammatory end-organ morbidities. Such life-threatening coronavirus disease 2019 (COVID-19) is heralded by virus-induced activation of mononuclear phagocytes (MPs; monocytes, macrophages, and dendritic cells). MPs play substantial roles in aberrant immune secretory activities affecting profound systemic inflammation and end organ malfunctions. All follow an abortive viral infection. To elucidate SARS-CoV-2-MP interactions we investigated transcriptomic and proteomic profiles of human monocyte-derived macrophages. While expression of the SARS-CoV-2 receptor, the angiotensin-converting enzyme 2, paralleled monocyte-macrophage differentiation it failed to affect productive viral infection. In contrast, simple macrophage viral exposure led to robust pro-inflammatory cytokine and chemokine expression but attenuated type I interferon (IFN) activity. Both paralleled dysregulation of innate immune signaling pathways specifically those linked to IFN. We conclude that the SARS-CoV-2-infected host mounts a robust innate immune response characterized by a pro-inflammatory storm heralding consequent end-organ tissue damage.

## Introduction

Severe acute respiratory syndrome coronavirus 2 (SARS-CoV-2), the causative agent of coronavirus disease 2019 (COVID-19), is an enveloped positive-stranded RNA virus belonging to the *Coronaviridae* family, *Betacoronaviruses* genus (Lu *et al*, 2020). COVID-19 has posed an unprecedented global threat to public health and in March of 2020, it was declared a pandemic by the World Health Organization (WHO) (World Health Organization, 2020). COVID-19 ranges from asymptomatic infection to mild pneumonia and, in its most severe form, progression to acute respiratory distress syndrome (ARDS). Such pulmonary compromise is associated with dyspnea and hypoxia that can progress to severely compromised lung dysfunction and multiorgan system failure and death (Gavriatopoulou *et al*, 2020). Disease mortality is linked to cytokine storm syndrome (CSS), heralded by innate immune activation with secretion of excessive pro-inflammatory cytokines (Ruan *et al*, 2020). Indeed, nearly 15% of reported COVID-19 disease cases progress to ARDS defined by widespread inflammatory-associated lung tissue damage and multiorgan failure (Guan *et al*, 2020) involving heart, liver, gastrointestinal tract, kidney, and brain (Salamanna *et al*, 2020). Viral persistence in the face of such end-organ disease is linked to cell expression of angiotensin-converting enzyme 2 (ACE2), the molecule that SARS-CoV-2 utilizes for receptor-mediated cell entry (Wan *et al*, 2020).

Mononuclear phagocytes (MPs; monocytes, macrophages, and dendritic cells) are the governors of innate immunity serving to contain microbial infection (Lugo-Villarino *et al*, 2019). Immediately following viral exposure, the process of intracellular microbial removal is initiated through recognition of viral pathogen-associated molecular patterns (PAMPs) by pattern recognition receptors (PRRs). This includes, but is not limited to, cytosolic retinoic acid-inducible gene I (RIG-I)-like receptors (RLRs) and extracellular and endosomal toll-like receptors (TLRs). During the initiation of virus-cell interactions, secretion of pro-inflammatory cytokines and chemokines initiate intracellular killing, antigen presentation, and mobilization of adaptive immunity. Factors that are engaged include, but are not limited to, interleukin-6 (IL-6), IL-1, tumor necrosis factor-alpha (TNF-α), and C-X-C motif chemokine ligand 10 (CXCL10) (Nikitina *et al*, 2018). Type I interferon (IFN) α and β responses occur in tandem to control both viral replication and dissemination (Swiecki & Colonna, 2011). During infection with SARS-CoV-2, CSS contributes to COVID-19-associated multiorgan failure (Huang *et al*, 2020; Xiong *et al*, 2020). In particular, infiltrating inflammatory MPs in lungs, heart, kidney, spleen, and lymph nodes are seen in post-mortem tissues of COVID-19-infected patients (Diao *et al*, 2021; Feng *et al*, 2020; Merad & Martin, 2020; Xu *et al*, 2020). Despite the critical role of MPs to clear the infection, these same cells underlie the pathobiology of COVID-19.

Mounting evidence confirms abortive MP infection by SARS-CoV-2 (Yang *et al*, 2020; Zheng *et al*, 2021). Nonetheless, such viral cell engagements are sufficient to induce activation and pro-inflammatory cytokine secretory responses. The nature of virus-MP interactions and their role in persistent viral infection and associated tissue pathologies remains enigmatic. To such ends, we pursued transcriptomic and proteomic analyses of immune cells following virus infection to identify modulations of host immune responses. We employed MPs to assess immune responses following SARS-CoV-2 cell engagements for cell activation using human immune response arrays and mass spectrometry-based label-free proteomic quantification methods. These techniques serve to define virus-induced innate immune responses linked to antiviral immunity. Based on these tests, we have uncovered the dysregulation of a spectrum of viral-induced responses related to IFN signaling pathways, complement activation, and linked adaptive immune responses. All provide unique insight into how the inflammatory response occurs as a consequence of SARS-CoV-2 infection. Virion persistence for up to 14 days after viral exposure despite abortive infection underlies the persistence of immune activation. Most importantly, the data provides a signature for virus-induced disease pathobiology, including COVID-19-associated CSS and multiorgan dysfunction. All affect the most severe disease morbidities and mortalities seen as a consequence of viral exposure, transmission, and dissemination.

## Materials and methods

### Isolation and cultivation of human monocytes

Human monocytes were obtained by leukapheresis from hepatitis B and HIV-1/2 seronegative donors and purified by counter-current centrifugal elutriation (Faradji *et al*, 1994). Monocytes were seeded in 6-well plates (3×10^6^ cells/well) in Dulbecco’s Modified Eagle’s Media (DMEM) containing 4.5 g/l glucose, L-glutamine, and sodium pyruvate, and supplemented with 10% heat-inactivated human serum, 50 μg/ml gentamicin, 10 μg/ml ciprofloxacin, and 1000 U/ml human macrophage colony-stimulating factor (M-CSF) to facilitate differentiation of monocytes into monocyte-derived macrophages (MDMs). Cells were incubated at 37°C with 5% CO_2,_ and the culture medium was half-exchanged with fresh medium every other day (Gendelman *et al*, 1992).

### Flow cytometry assays

Human monocytes were evaluated by flow cytometry for levels of SARS-CoV-2 cell entry receptor ACE2 and monocyte-macrophage phenotypic surface markers, CD14 and CD16, during macrophage differentiation with or without captopril (Sigma-Aldrich, C4042). Captopril was added to the culture medium to increase ACE2 cell expression. On days 0, 1, 3, 5, and 7 during differentiation, monocytes-macrophages were stained with fluorescently-conjugated antibodies to detect human ACE2 (APC, LSBio, LS-C275129, polyclonal), CD14 (Alexa Fluor 488, eBioscience, clone 61D3), and CD16 (PE, eBioscience, eBioCB16, clone CB16), and with isotype-matched antibodies serving as negative controls. Stained cells were examined with an LSR II flow cytometer (BD Biosciences) and analyzed using BD FACSDiva software.

### SARS-CoV-2 infection

Experiments involving SARS-CoV-2 were performed in the University of Nebraska Medical Center (UNMC) biosafety level 3 (BSL-3) core facility and approved by UNMC Institutional Biosafety Committee (IBC) (protocol number 20-05-027-BL3). The SARS-CoV-2 strain used in this study was isolate USA-WI1/2020 (BEI, NR-52384). The virus was passaged on Vero.STAT1 knockout (KO) cells (ATCC, CCL-81-VHG) and titer was determined by plaque assay in Vero E6 cells (ATCC, CRL-1586) (Mendoza *et al*, 2020). On day 5 following cell cultivation, MDMs were exposed to the WI1/2020 viral strain at a multiplicity of infection (MOI) of 0.01. Exposure was maintained with or without captopril, and the virus-cell mixtures incubated at 37°C in 5% CO_2_ with shaking at 15 minute intervals for 1 hour. This was followed by incubation with the virus for an additional 3 hours. At the termination of viral cell incubation, the virus inoculum was removed, and cells were washed 3 times with phosphate-buffered saline (PBS). Mock-challenged cells were treated with a culture medium alone. Culture supernatants were collected at defined time points then used for viral quantitative reverse transcription-polymerase chain reaction (RT-qPCR) and type I IFN activity assays. Vero.STAT1 KO cells were maintained for study based on their high susceptibility to virus infection due to lack of Signal Transducer and Activator of Transcription 1 (STAT1) protein required for cellular antiviral responses (Durbin *et al*, 1996). Thus, the SARS-CoV-2 replication kinetics in MDMs were compared against viral infection in Vero.STAT1 KO cells that were monitored over 5 days post-viral exposure.

### RT-qPCR assay

Culture supernatants were collected on days 1, 3, 5, 7, and 10 following viral exposure. Viral RNA was extracted using QIAamp Viral RNA Mini Kit (Qiagen, 52906). SARS-CoV-2 genome equivalents were quantified in culture supernatant by RT-qPCR using 2019-nCoV CDC probe and Primer Kit for SARS-CoV-2 (Biosearch Technologies, KIT-nCoV-PP1-1000). Forward primer: 5’ GACCCCAAAATCAGCGAAAT 3’, Reverse primer: 5’ TCTGGTTACTGCCAGTTGAATCTG 3’, and Probe: 5’ FAM-ACCCCGCATTACGTTTGGTGGACC-BHQ-1 3’. RT-qPCR was performed using Taqman Fast Virus 1-step Master Mix (Thermo Fisher Scientific, 4444434) in StepOne Plus real-time PCR thermocycler (Applied Biosystems) using the following cycling conditions: 50°C for 10 minutes, 95°C for 3 minutes, and 40 cycles of 95°C for 15 seconds, followed by 60°C for 1 minute. The SARS-CoV-2 genome equivalent copies were calculated using control RNA from heat-inactivated SARS-CoV-2 USA-WA1/2020 (BEI, NR-52347).

### Transmission electron microscopy (TEM)

For negative staining analysis of viral particles used for cell infection experiment, purified SARS-CoV-2 viruses were de-activated and fixed in 2% glutaraldehyde and 2% paraformaldehyde (PFA) in 0.1 M Sorenson’s phosphate buffer for 1 hour at room temperature. Briefly, for negative staining, a drop of the virus sample in a fixative solution was placed onto a formvar and carbon-coated grid for 30-40 seconds. After removing excess sample solution with filter paper, the grid was placed with sample side down on a drop of 1% phosphotungstic acid (PTA) in water, stained for 30 seconds, then excess PTA solution was blotted with filter paper. Stained samples were examined and imaged using a Hitachi H7500 TEM (Hitachi High-Tech GLOBAL) and a bottom-mount AMT camera (AMT Imaging). For ultrastructural analysis, mock and SARS-CoV-2-challenged MDMs sampled at 1, 3, 5 and 14 days post-viral inoculation were washed 2 times with PBS and fixed in a solution of 2% glutaraldehyde and 2% PFA in 0.1 M Sorenson’s phosphate buffer for 24 hours at 4°C which have been washed 3 times with PBS to clear excess fixative solution. TEM analysis was performed as previously described (Mukadam *et al*, 2020) in samples post-fixed in a 1% aqueous solution of osmium tetroxide for 30 minutes that were dehydrated in 50, 70, 90, 95, and 100% graded ethanol. Spurr’s resin was used as embedding medium after solvent transition with ethanol and Spurr’s resin (50:50 ethanol:resin, followed by twice immersion in 100% Spurr’s resin for 2-3 hours for each solution), and embedded samples were cured at 60 - 65°C for 24 hours. Ultrathin sections (100 nm) were cut with Leica UC7 ultramicrotome, placed on 200-mesh copper grids, followed by staining with 2% uranyl acetate and Reynold’s lead citrate, and examined with a Hitachi H7500 TEM at 80 kV. Images were acquired digitally with an AMT digital imaging system.

### Transcriptomic analyses

SARS-CoV-2-challenged MDMs were collected on days 1, 3, and 5 post-viral exposure. Total RNA was isolated using RNeasy Mini Kit (Qiagen, 74104), and cDNA was generated utilizing RevertAid First Strand cDNA synthesis kit (Thermo Fisher Scientific, K1622) followed by amplification and quantification using RT^2^ Profiler Human Innate and Adaptive Immune Response 96-well Array (Qiagen, 330231) with RT^2^ SYBR Green ROX qPCR Mastermix (Qiagen, 330523).

The qPCR cycling conditions were 95°C for 10 minutes for 1 cycle, followed by 40 cycles of 95°C for 15 seconds and 60°C for 1 minute using Eppendorf Mastercycler ep realplex 2S. Fold changes were determined via Qiagen’s RT^2^ Profiler analysis software (version 3.5). Ingenuity Pathway Analysis (IPA) (Qiagen) was used to identify the pathways and networks affected post-viral exposure. Functional and pathway enrichment analyses of screened genes in SARS-CoV-2-challenged MDMs were compared to mock-challenged MDMs at different time intervals following viral exposure. Gene ontology (GO) annotation was conducted using the GO Resource database (http://geneontology.org). The Kyoto Encyclopedia of Genes and Genomes (KEGG) pathway enrichment analysis was conducted utilizing DAVID (http://david.abcc.ncfcrf.gov/) (Huang da *et al*, 2009), an online tool providing a comprehensive set of functional annotations for understanding biological meaning in large list of genes. The Reactome gene set enrichment analysis (GSEA) was conducted using ReactomeFIViz (https://reactome.org/tools/reactome-fiviz) (Wu *et al*, 2014), a Cytoscape application for pathway and network-based data analysis. The Search Tool for the Retrieval of Interacting Genes/Proteins (STRING) local network cluster enrichment analysis was conducted using STRING database (http://string-db.org), which provides a critical assessment and integration of protein-protein interaction (PPI), including direct (physical) and indirect (functional) associations in a given organism (Szklarczyk *et al*, 2019).

### Measures of IFN activity

MDBK cells (ATCC, CCL-22) were cultured in Eagle’s Minimum Essential Medium (EMEM) containing 10% heat-inactivated fetal bovine serum (FBS) and 50 µg/ml gentamicin (Rubinstein *et al*, 1981). Vesicular stomatitis virus (VSV Indiana laboratory strain) (V-520-001-522, ATCC, VR-1238) was passaged on Vero cells (ATCC, CCL-81), and viral titer was determined using the plaque assay in Vero cells. MDBK cells bind and respond to human type I IFNs, α and β, but not IFN-gamma (IFN-γ) (Rubinstein *et al*., 1981). Culture supernatants were collected from virus-exposed MDMs on days 1, 3, and 5 following infection with SARS-CoV-2 at MOI of 0.01, then assessed for IFN-α/β activity by protection against the VSV-induced cytopathicity measured in MDBK cells (Rubinstein *et al*., 1981). Polyinosinic-polycytidylic acid (poly(I:C); Sigma-Aldrich, P9582) served as a positive control for IFN induction via TLR3 engagement (Field *et al*, 1967). Recombinant human IFN-α (PBL Assay Science, 11200-2) was used as assay standard. We also investigated IFN activity upon the challenge of Teflon flask-suspended monocytes with SARS-CoV-2 (MOI=0.01), before and after treatment of cells with 100 µg/ml poly(I:C).

### Western blot analysis

At days 1, 3, and 5 after infection, MDMs were collected, and total protein was extracted using lysis buffer containing 0.1% SDS, 100 mM Tris-HCL, 150 mM NaCl, 1 mM EDTA, 1% Triton X-100, supplemented with protease and phosphatase inhibitor cocktail (Sigma-Aldrich, PPC1010). Protein concentration was determined utilizing Pierce 660 Protein Assay kit (Thermo Fisher Scientific, 22662) with ionic detergent compatibility reagent (Thermo Fisher Scientific, 22663) following the manufacturer’s instructions. Protein lysates (25 μg) were resolved by SDS-PAGE and transferred to Immobilon-P PVDF membrane (Sigma-Aldrich, IPVH00010). Membranes were blocked in 5% nonfat milk in TBST buffer at room temperature for 1 hour, followed by incubation with primary antibodies to IFN-α (1:500, Thermo Fisher Scientific, MA5-37518), IFN-β (1:1000, Abcam, ab85803), and β-actin (1:3000, Sigma-Aldrich, A3854) at 4°C overnight, followed by 1 hour incubation in 3% nonfat milk in Tris-buffered saline with 0.1% Tween 20 detergent (TBST) buffer with horseradish peroxidase-conjugated anti-rabbit (1:2000, R&D Systems, HAF008) or mouse (1:2000, R&D Systems, HAF018) secondary antibody. Immunoreactive bands were detected using SuperSignal West Pico Chemiluminescent substrate (Thermo Fisher Scientific, 34080), and images were captured using an iBright CL750 Imager (Thermo Fisher Scientific). Immunoblots were quantified using ImageJ software (NIH) relative to β-actin expression.

### Proteomic analysis

On days 1, 3, and 5 post-infection, MDMs were collected, lysed with 2% SDS in 100 mM Tris-HCL and 100 mM dithiothreitol, pH 7.6, and supplemented with protease and phosphatase inhibitors. Protein concentration was determined using Pierce 660 Protein Assay kit with ionic detergent compatibility reagent according to the manufacturer’s instructions. Afterward, samples were processed as previously described (Arainga *et al*, 2015) using filter-aided sample preparation (FASP, Pall Life Sciences, OD010C34) for digestion of 50 μg per sample. Following overnight digestion, samples were cleaned using the Oasis MCX column (Waters, 186000252) and C18 Zip-Tips (Sigma-Aldrich, ZTC18M960). Cleaned peptides were quantitated using NanoDrop2000 at 205 nm. Following resuspension in 0.1% formic acid, 2 μg of the sample was used for label-free quantification (LFQ) in UNMC Mass Spectrometry and Proteomics Core Facility as previously described (Gao *et al*, 2020). Proteins identified by mass spectrometry were quantified to determine differentially expressed proteins between mock and SARS-CoV-2-challenged MDMs at different time points post-infection. Statistical analysis for proteomic data was performed using ANOVA, and false discovery rate (FDR) for pathway analysis was controlled using the Benjamini-Hochberg (BH) method (Benjamini & Hochberg, 1995). A protein was considered to be differentially expressed if p value ≤ 0.05 and the absolute value of fold change ≥ 2. Next, functional and pathway enrichment analysis of differentially expressed proteins in SARS-CoV-2-challenged MDMs compared to mock-challenged MDMs was conducted at different times after virus challenge. Gene enrichment analysis to identify immune system processes affected after virus exposure was performed using Cytoscape in conjunction with the plug-in ClueGO (Bindea *et al*, 2009). GO annotation, KEGG, Reactome GSEA, and STRING analyses were conducted using GO Resource, DAVID, ReactomeFIViz, and STRING databases, respectively.

## Statistical analysis

Results are presented as the mean ± standard error of the mean (SEM). Student’s t-test or one-way ANOVA followed by Tukey’s multiple comparison test was used to analyze differences in the mean values between groups. P values ≤ 0.05 were considered statistically significant. Statistical analysis was performed using GraphPad Prism 9.1.0 software (GraphPad Software, San Diego, CA).

## Results

### ACE2 receptor expression during human monocyte-macrophage maturation

ACE inhibitors were used to upregulate the SARS-CoV-2 receptor ACE2 and potentiate cell entry (Ferrario *et al*, 2005; Pedrosa *et al*, 2021; Soler *et al*, 2009). As ACE inhibitors are frequently prescribed to treat hypertension, we posit that they can increase susceptibility to SARS-CoV-2. Therefore, we measured ACE2 expression on freshly isolated monocytes throughout cell isolation and differentiation. Days 1, 3, 5, and 7 were evaluated for cells cultured with or without captopril, an ACE inhibitor which is thought to increases ACE2 expression. In these studies, ACE2 expression by monocytes was found to peak by day 5 after initiation of cell differentiation then decreased after that (**Figure 1A**), which corresponds to the susceptibility of infection of monocytes-macrophages by other viruses (Gendelman *et al*., 1992; Lum *et al*, 2018; Schmidtmayerova *et al*, 1996). On day 5, ACE2 expression increased by 23% in captopril-treated cells. In parallel, the expression of phenotypic monocyte-macrophage surface markers, CD14 and CD16, were not changed (**Figure 1A**). Based on these observations, we challenged MDMs with SARS-CoV-2 on day 5 after start of cell differentiation and assessed whether captopril affects susceptibility to infection.

**Figure 1.**
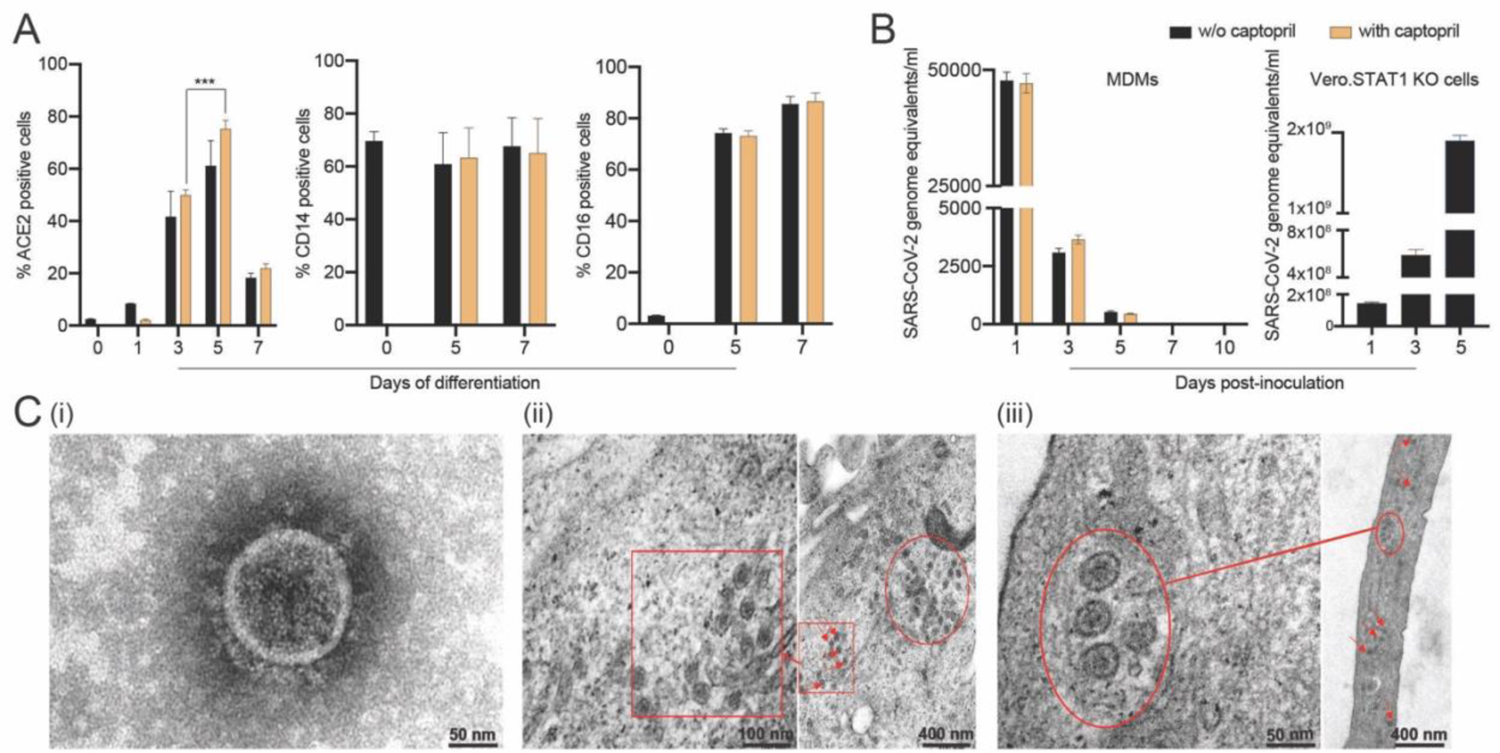
SARS-CoV-2 abortive infection of human MDMs. (A) Expression of SARS-CoV-2 cell entry receptor ACE2 and phenotypic surface markers CD14 and CD16, during differentiation of monocytes into macrophages, was analyzed by flow cytometry in absence or presence of captopril. (B) SARS-CoV-2 presence in MDMs. SARS-CoV-2 (MOI=0.01) was used to infect MDMs (with or without captopril). The number of virus genome equivalents per ml was measured in the culture supernatant by RT-qPCR. Vero.STAT1 KO cells served as a positive control. (C) Transmission electron micrographs of SARS-CoV-2-challenged MDMs. (C.i) A viral particle from a pool of SARS-CoV-2 used for the virus challenges. SARS-CoV-2-challenged MDMs (C.ii: day 5, and C.iii: day 14 after viral exposure). Red arrows, circles, and boxes demonstrate clusters of viral particles within the macrophage cytoplasm. All experiments were done at least twice with representative images depicted here. (**A** and **B**) Data are represented as mean ± SEM (n=3-6). Statistical significance between groups was determined using unpaired Student’s t-test or one-way ANOVA, and p < 0.05 was considered significant (*** p < 0.001). Abbreviations: w/o: without. Scale bars: 50 nm (i and left panel of iii), 100 nm (left panel of ii), and 400 nm (right panels of ii and iii).

### Abortive infection of MDMs is established following SARS-CoV-2 exposure

To investigate whether human MDMs are susceptible to SARS-CoV-2 infection, monocytes were isolated from healthy donors and differentiated in the presence or absence of captopril and infected at day 5 after cell culture. The kinetic growth of SARS-CoV-2 in MDMs was determined from culture supernatant tests of challenged cells over 10 days post-infection. In all studied groups (captopril-treated and controls), the SARS-CoV-2 genome equivalent copies in the supernatant significantly decreased during 5 days of viral exposure, becoming undetectable after that (**Figure 1B**). Additionally, the number of genome copies did not change significantly with or without captopril. These results demonstrated that SARS-CoV-2 infection of human MDMs was abortive and differentiated monocytes could not generate progeny virus in contrast to productive infection of SARS-CoV-2 in Vero.STAT1 KO cells (**Figure 1B**). Our data also indicated that captopril did not alter the susceptibility of MDMs to SARS-CoV-2 infection.

### Ultrastructural features of SARS-CoV-2-challenged MDMs

Viral particles and mock and SARS-CoV-2-challenged MDMs were fixed at different time points after viral exposure and then examined by TEM to determine ultrastructural changes in the virus-challenged cells (**Figure 1C.i-iv**). Negative stain TEM shows the ultrastructure of a viral particle from a pool of SARS-CoV-2 used for the virus challenges (**Figure 1C.i**). By day 5 after exposure, a cluster of viral particles was shown in the cytoplasm of challenged cells **(Figures 1C.ii)**. Surprisingly, while no viral genome copies were detected after 5 days (**Figure 1B**), mature virions were observed 14 days after initial exposure (**Figure 1C.iii**). These underly the persistence of virus in macrophages despite evidence of abortive infection.

### Abortive SARS-CoV-2 infection induces pro-inflammatory factors, but not IFN

To investigate alterations in macrophage-mediated innate immune responses after SARS-CoV-2 exposure, a total of 84 key genes were examined. Transcriptional changes linked to immunity were screened using RT^2^ Profiler Human Innate and Adaptive Immune Response Array. Expression of immune response genes in SARS-CoV-2-challenged MDMs were investigated against control “mock-challenged cells” on days 1, 3, and 5 (**Supplementary Figure 1**). On day 1, more than two-fold increase in expression of pro-inflammatory cytokine and chemokine genes were detected, including *IL-6, TNF-α, IL-1α, IL17A, IL8*, colony-stimulating factor 2 (*CSF2*), chemokine (C-C motif) ligand 2 (*CCL2*), and *CCL5*. Interestingly, on day 3, gene expression of the node-like receptor family pyrin domain containing 3 (*NLRP3*) and *IL-1β* increased by 8.84- and 5.64-fold, respectively. On day 5, expression for these pro-inflammatory cytokines and chemokines were at baseline or decreased with the exception of *IL-6* and *CCL2*. Gene expression of anti-inflammatory cytokine *IL-10* was elevated on days 1 and 5 supporting the attenuation of pro-inflammatory responses (Lu *et al*, 2021). Importantly, expression of IFN-linked genes *IFN-α1, IFN-β1, IFN-γ*, IFN regulatory factor 7 (*IRF7*), and tyrosine kinase 2 (*TYK2*) genes increased on day 1 but returned to near baseline by days 3 and 5. These data demonstrate a complex pro- and anti-inflammatory network interactions that are sustained to day 5 with attenuated IFN responses following SARS-CoV-2 exposure (**Figure 2A**).

**Figure 2.**
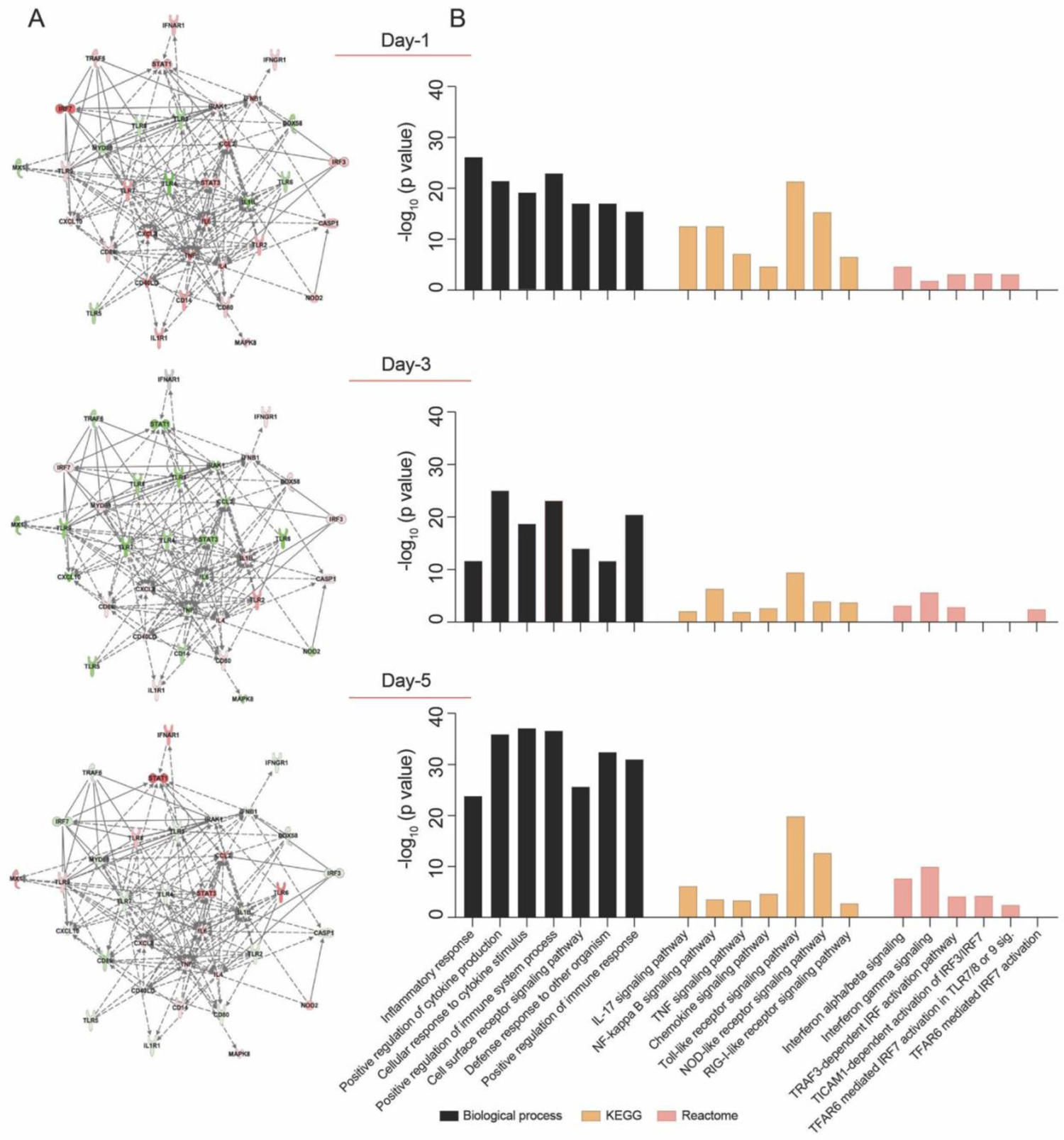
Transcriptomic profile of SARS-CoV-2-challenged MDMs. Expression of 84 genes specific for human innate and adaptive immune responses were screened in SARS-CoV-2-challenged MDMs compared to mock-challenged MDMs at different times post-challenge, using RT^2^ Profiler Human Innate and Adaptive Immune Response 96-well Array. Fold changes in the gene expression were determined via Qiagen’s RT^2^ Profiler analysis software (n=4). (A) IPA was performed with upregulated or downregulated genes to identify putative network interactions involved in SARS-CoV2-MDM interactions. Upregulated genes are shaded red with darker shade indicating higher upregulation, while green shades denote downregulation in gene expression. Solid grey lines indicate direct interactions, while dotted grey lines correspond to indirect interactions. (B) Functional and pathway enrichment analysis of transcriptomic dataset was performed using GO-term, KEGG, and Reactome analyses at different time points post-infection. Different immune and inflammation-related biological processes and pathways affected in SARS-CoV-2-challenged MDMs were plotted as a bar chart compared to mock-challenged MDMs.

### Functional and pathway enrichment of immune regulated genes

SARS-CoV-2 infection may induce dynamic changes of immune-based gene expression in specific cellular biological processes and pathways in virus-challenged cells. To assess these changes, functional and pathway enrichment analyses were performed on immune-regulated genes screened in SARS-CoV-2-challenged MDMs compared to controls at days 1, 3, and 5 (**Supplementary file 1**). Multiple immune processes were enriched upon viral exposure, including defense response to other organism, positive regulation of immune system processes, cell surface receptor signaling pathways, and positive regulation of immune responses (**Figure 2B**). All indicated they comprised activated immune responses in the virus-challenged MDMs. In addition, a series of inflammation-related processes were enriched, including inflammatory response, positive regulation of cytokine production, and cellular response to cytokine stimulus (**Figure 2B**), highlighting inflammation responses induced after viral exposure. Similarly, different immune and inflammation-related molecular functions such as cytokine activity, chemokine activity, complement component C1q binding, type I IFN receptor binding, and IL-1 receptor binding were enriched after virus exposure (**Supplementary file 1**). Furthermore, KEGG pathway analysis showed enrichment of TLR, NOD-like receptor, and RLR signaling pathways (**Figure 2B**), suggesting the activation of PRRs on MDMs to recognize viral PAMPs initiating innate immune responses. Moreover, KEGG pathway analysis depicted enrichment of IL-17, NF-κB, TNF, and chemokine signaling pathways (**Figure 2B**), highlighting different inflammatory pathways induced in the SARS-CoV-2-challenged MDMs. Interestingly, Reactome analysis revealed enrichment of multiple IFN-related signaling pathways including IFN-α/β signaling, IFN-γ signaling, TICAM1-dependent activation of IRF3/IRF7, TRAF3-dependent IRF activation pathway, and TRAF6 mediated IRF7 activation (**Figure 2B**). In addition, immune and inflammation-mediated molecular functions were enriched in virus-challenged cells (**Supplementary file 1**). The interacting proteins involved in the screened innate and adaptive immune-related genes were also identified using STRING analysis of their interaction networks (**Supplementary file 1**).

### SARS-CoV-2 monocyte-macrophage engagement and failed induction of IFN activity

To determine whether exposure to the SARS-CoV-2 can trigger antiviral IFN activities in human macrophages, culture supernatants were collected at different times post-viral exposure and assayed in VSV-challenged MDBK cells. Notably, no type I IFN activity was observed in SARS-CoV-2-challenged MDM cultures at different time intervals following viral challenge (**Figure 3A).** These data stand in contrast to the transcriptomic results, which displayed increases in IFN pathway-linked genes. Furthermore, no additive or synergistic IFN activity was detected in control and SARS-CoV-2-challenged monocytes treated with poly(I:C) (**Figure 3B**). Taken together, the data demonstrate that SARS-CoV-2-MDM interactions do not affect IFN activities. To affirm this, we evaluated the protein expression level of IFN-α and IFN-β in SARS-CoV-2 exposed MDMs at days 1, 3, and 5 post-exposure. Similar to IFN activity, no IFN-α or IFN-β proteins were produced in the SARS-CoV-2-challenged MDMs (**Figure 3C**). Altogether, our data demonstrate that SARS-CoV-2 exposure of MDM does not trigger IFN activity or protein production in human monocytes-macrophages.

**Figure 3.**
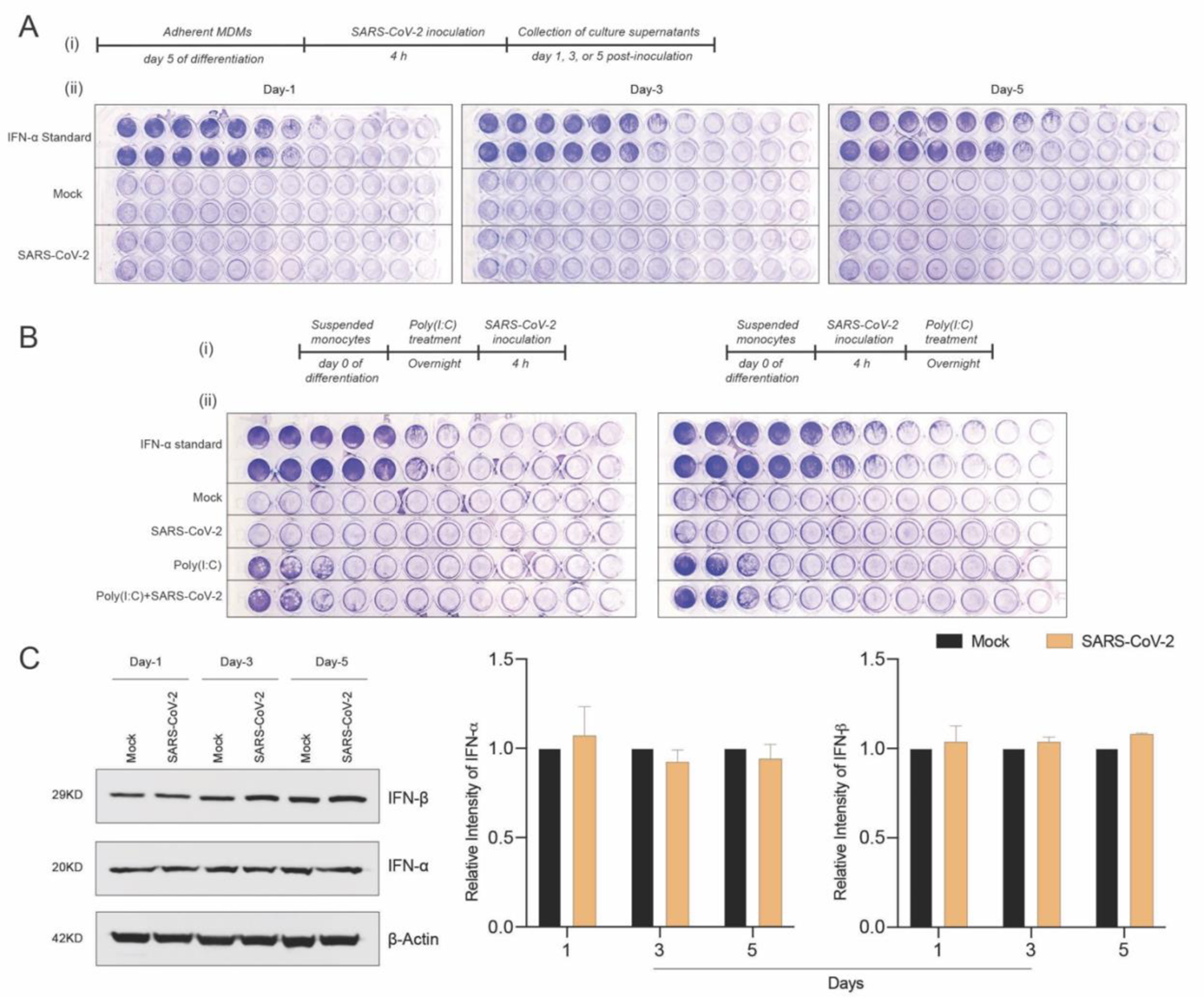
SARS-CoV-2 does not induce IFN activity in monocytes-macrophages. (A) MDMs were infected with SARS-CoV-2 (MOI=0.01) and the culture supernatants were collected at days 1, 3, and 5 post-infection. The collected culture supernatants were used to assess IFN activity in VSV-challenged bovine MDBK cells. Recombinant human IFN-α was used as standard. (A.i) Experimental timeline. (A.ii) Representative images of IFN activity assay plates. (B) No additive or synergistic responses in poly(I:C)-induced IFN activity between control and SARS-CoV-2-challenged monocytes. Suspended monocytes were treated with 100 μg poly(I:C) overnight before or after the infection with SARS-CoV-2 (MOI=0.01) for 4 h. (B.i) Experimental timeline. (B.ii) Representative images of IFN activity assay plates. (C) Western blot analysis was performed to determine expression of IFN-α and IFN-β in cell lysates at different times after viral exposure. Representative immunoblot and densitometric quantification are shown. All experiments were done at least twice, and one representative image is shown. (C) Data represent mean ± SEM (n=3). Statistical significance between the groups was determined with unpaired Student’s t-test and p < 0.05 was considered significant.

### Proteomic profiles of SARS-CoV-2-challenged MDMs

In attempts to elucidate the mechanisms of the MDM response against the SARS-CoV-2 challenge, we obtained the proteome profiles from lysates of viral exposed macrophages and compared those to non-exposed counterparts. On defined days after viral exposure, the expression of more than 4,000 proteins were identified and quantified using differential proteomic tests. Amongst total identified proteins, 1776, 1372, and 2448 proteins were significantly differentially expressed (p ≤ 0.05 and fold change ≥ 2), on days 1, 3, and 5, respectively, after viral challenge (**Supplementary file 2**). Volcano plots depicting differentially expressed proteins in SARS-CoV-2-challenged MDMs compared to controls are shown in **Supplementary Figure 2**. To obtain a detailed understanding of changes in the MDM proteomic profile after viral challenge, we performed functional and pathway enrichment analyses of differentially regulated proteins in virus-challenged MDMs at different times. Amongst total proteins altered on day 1, only 5% were involved in negative regulation of inflammatory response to an antigenic stimulus. In contrast, on days 3 and 5, 38.1% and 23.53% of altered proteins, respectively, belonged to myeloid cell activation involved in immune responses (**Figure 4A**). Immune processes of each function indicated in pie charts include, but are not limited to, activation of the innate immune response, myeloid cell activation, innate immune response activating signal transduction, and innate immune response activating cell surface receptor signaling pathway (**Figure 4A** and **Supplementary file 3**). All indicated activated immune and inflammatory responses in MDMs upon viral challenge. Additionally, viral exposure induced changes in biological processes related to the endoplasmic reticulum (ER) and mRNA on days 1 and 3 after challenge. With positive fold changes, the enriched biological processes included protein localization to ER, protein targeting to ER, regulation of mRNA metabolic process, nuclear-transcribed mRNA catabolic process, and mRNA splicing via spliceosome (**Supplementary file 4**). Similarly, on days 1 and 3 after viral challenge, Reactome analysis depicted enrichment of processes related to RNA processing, with positive fold change, including rRNA processing in the nucleus and cytosol and mRNA splicing (**Supplementary file 4**). Moreover, KEGG pathway analysis showed an enrichment of ribosome function as a pathway of genetic information processing and translation on day 1. Similarly, the molecular function of a structural constituent of the ribosome was enriched with positive fold change on day 1 (**Supplementary file 4**). Interestingly, GO-term analysis showed robust enrichment of different immune responses such as complement activation, humoral immune response, adaptive immune response, and regulation of inflammatory response with negative fold changes suggesting dysregulated immune responses to SARS-CoV-2 in MDMs (**Figure 4B**). Importantly, STRING analysis of PPIs showed enrichment of IFN-related interaction networks such as “IFN signaling and positive regulation of RIG-I signaling pathway” and “IFN-α/β signaling and IFN-γ signaling”, with negative fold changes, suggesting dysregulated IFN responses induced by SARS-CoV-2 (**Figure 4C** and **Supplementary file 4**). Overall, these results support transcriptomic, IFN activity, and protein evaluations that demonstrated attenuated IFN responses in SARS-CoV-2-challenged MDMs.

**Figure 4.**
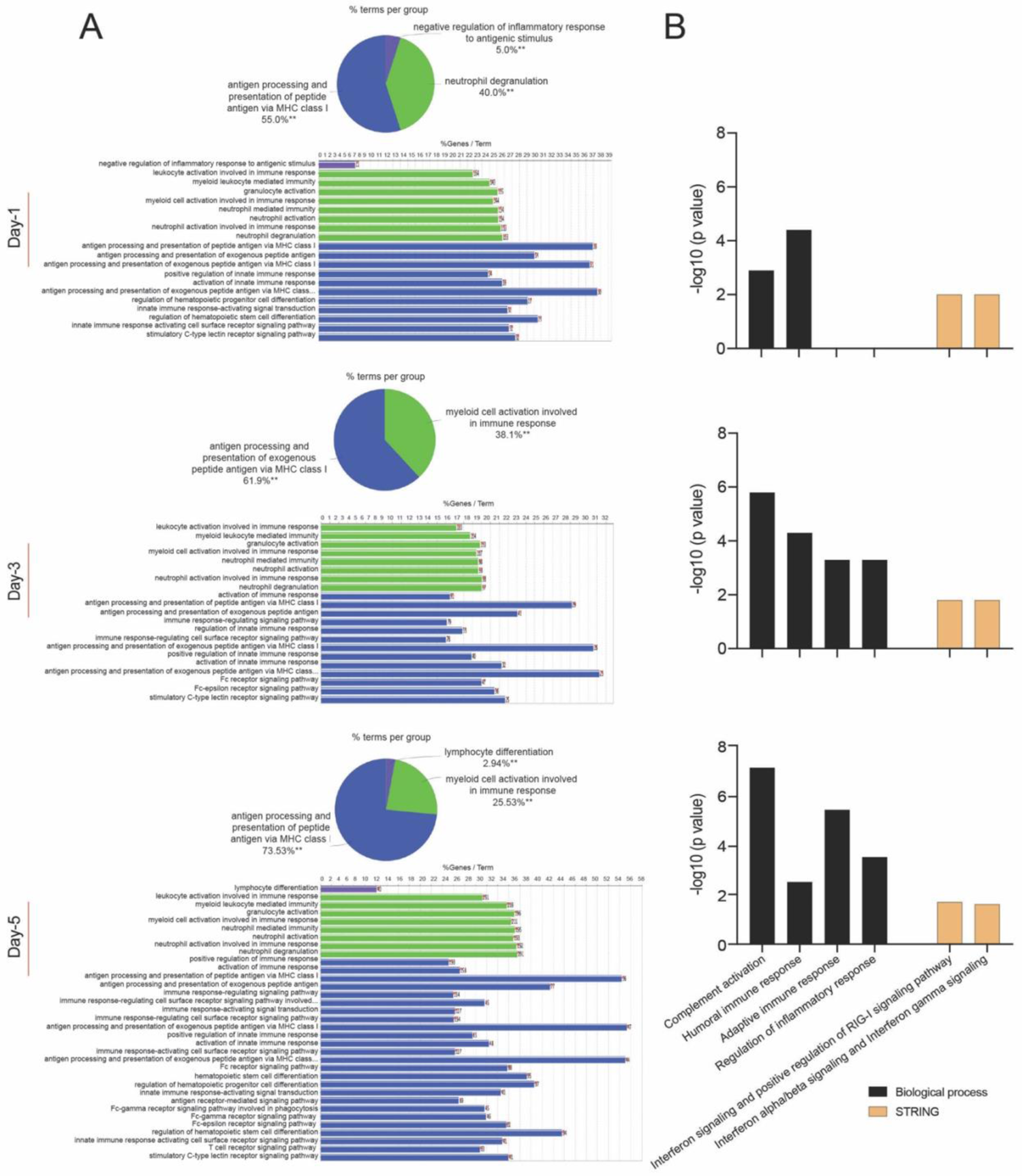
Proteomic profile of SARS-CoV-2-challenged MDMs. (A) Gene enrichment analysis was performed using Cytoscape in conjunction with the plug-in ClueGO at different times after viral exposure. Pie charts representing the distribution of identified differentially expressed proteins according to observed immune processes and bar charts demonstrating the specific processes that correspond to their classification. The same color key used in the pie charts applied in the bar charts. (B) Functional and pathway enrichment analyses of proteomic dataset were performed using GO annotation, KEGG, Reactome GSEA, and STRING analyses at different time points post-infection. Different immune-related biological processes and PPI networks affected in SARS-CoV-2-challenged MDMs were plotted as a bar chart compared to mock-challenged MDMs.

## Discussion

MPs are the first line of host defense against viral infection (Nikitina *et al*., 2018). These sentinel cells interact with SARS-CoV-2 to define COVID-19 pathogenesis. That includes, but is not limited to, CSS, ARDS, and multiorgan dysfunction (Guan *et al*., 2020). As a first line defense mechanism, the initial virus-MP interaction following the entry of the pathogenic virus into host cells represents a critical determinant for infectivity and pathogenesis (Li, 2016; Perlman & Netland, 2009; Shang *et al*, 2020). To this end, we have better defined such virus-host cell interactions and determined relationships between ACE2 receptor expression and monocyte-macrophage differentiation during abortive SARS-CoV-2 infection. Like other viral infections, we reasoned that monocytes would respond with an elevated expression of pro-inflammatory molecules and antiviral responses. Indeed, this has already been reported for influenza, Chikungunya, herpes, Zika, and lentiviral infections that include human immunodeficiency virus type one (HIV-1) and herpes viruses (Coates *et al*, 2018; Her *et al*, 2010; Krzyzowska *et al*, 2013; Lum *et al*., 2018; Schmidtmayerova *et al*., 1996). Each elicits profound inflammation during early viral infection. However, unlike other known viral infections, SARS-CoV-2 infection is abortive and balances pro- and anti-inflammatory processes with a unique cell signature. The resulting signature defines the pathways towards the major cause for morbidity and mortality in COVID-19: the CSS leading to ARDS and other multi-organ failures. The specific lack of induction of IFN also serves to define a specific monocyte-macrophage-virus phenotype during both early and progressive infection (Dalskov *et al*, 2020; Yang *et al*., 2020).

Herein, we show increased expression of viral receptor ACE2 during monocyte-macrophage differentiation. Interestingly, further induction of ACE2 expression by captopril, a well-known ACE inhibitor, was observed. Captopril induces a reduction of angiotensin II, which increases ACE2 expression and activity. This occurs through angiotensin type 1 receptor-dependent ACE2 internalization followed by lysosomal degradation (Deshotels *et al*, 2014). However, the observed upregulation of ACE2 expression failed to affect infection with SARS-CoV-2. Despite infection of MDMs with SARS-CoV-2 when ACE2 receptor expression peaked, the infection remained abortive in agreement with prior reports (Boumaza *et al*, 2021; Yang *et al*., 2020; Zheng *et al*., 2021). The inability of captopril to increase the susceptibility of MDM infection with SARS-CoV-2 is in accordance with a recent study indicating that treatment of human alveolar type-II pneumocytes with captopril induced upregulation of ACE2 expression and counteracted drug-induced reduction of SARS-CoV-2 spike protein entry (Pedrosa *et al*., 2021). This occurred through inhibition of A disintegrin and metalloprotease 17 (ADAM17), which has been shown to play an essential role in ACE2 shedding and viral entry into the cells (Lambert *et al*, 2005). Nonetheless, viral particles were present inside cytoplasmic structures of macrophages up to two weeks after the viral challenge underscoring any continuity of immune responses.

SARS-CoV-2 abortive infection likely results from intracellular mechanisms induced upon MP activation. Human coronaviruses can infect human peripheral blood mononuclear cells leading to cell activation and aberrant production of pro-inflammatory mediators with increased chemoattraction (Channappanavar & Perlman, 2017; Yilla *et al*, 2005; Zhou *et al*, 2014). In particular, virus-exposed monocytes-macrophages can serve as perpetrators for virus-induced inflammatory responses within different body organs, as seen by the abundance of pro-inflammatory macrophages in bronchoalveolar lavage fluid obtained from severe COVID-19 cases (Liao *et al*, 2020). The MP pro-inflammatory factors can contribute to local tissue inflammation and systemic inflammatory responses that characterize cytokine storm (Nikitina *et al*., 2018). In the current study, the transcriptomic profile of SARS-CoV-2-challenged MDMs demonstrated increased mRNA expression of multiple pro-inflammatory molecules, including *IL-6, TNF-α, IL-1α, IL17A, IL8, CSF2, CCL2, CCL5, NLRP3,* and *IL-1β* upon the virus exposure. Our data confirmed previous reports which showed excessive production of pro-inflammatory cytokines and chemokines in the SARS-CoV-2-exposed MPs (Boumaza *et al*., 2021; Yang *et al*., 2020; Zheng *et al*., 2021). In parallel, our data displayed enrichment of different inflammation-related pathways such as IL-17, NF-κB, and TNF signaling pathways, following viral exposure. IL-17 is known for its pivotal role in inducing and mediating pro-inflammatory responses and its involvement in different inflammatory autoimmune diseases (Kuwabara *et al*, 2017). NF-κB serves as a central mediator of inflammation since the DNA binding site for NF-κB was found in the promoter regions of multiple pro-inflammatory molecules (Hayden & Ghosh, 2004). Thus, the activation of NF-κB with other pro-inflammatory transcription factors leads to the transcription of several pro-inflammatory molecules such as IL-1β, inducible nitric oxide synthase (iNOS), and TNF-α (Dasgupta *et al*, 2003; Liu *et al*, 2002). In addition, the transcriptome of SARS-CoV-2-challenged MDMs showed enrichment of multiple immune-related biological processes, molecular functions, and signaling pathways such as positive regulation of immune response, cell surface receptor signaling pathway, cytokine activity, chemokine activity, TLR signaling pathway, and NOD-like receptor signaling pathway, demonstrating cell activation of MDMs after viral exposure. Notably, the proteome of virus-challenged macrophages revealed the dysregulation of different immune processes such as regulation of inflammatory response, complement activation, and linked adaptive and humoral immune responses. Overall, our findings demonstrate MDM activation by SARS-CoV-2 is associated with exaggerated inflammatory responses and dysregulated immune activities.

Significantly, SARS-CoV-2 induced an attenuated MP IFN response, as the transcript expression of type I (IFN-α1 and IFN-β1) and type II (IFN-γ) IFNs in the virus-challenged cells increased early after exposure but returned to or below normal levels at later times. Our findings are consistent with previous reports depicting a dysregulated IFN response in SARS-CoV-2-challenged MDMs and alveolar macrophages (Dalskov *et al*., 2020; Yang *et al*., 2020). This attenuated IFN response provides a possible mechanism for delayed viral clearance from cells up to two weeks after inoculation. Similarly, our study showed the inability of SARS-CoV-2 to induce IFN activity or production in MDMs, as illustrated by the failure of culture supernatants of infected cells to provide protection against the cytopathic effects induced by VSV in MDBK cells as well as unchanged levels of IFN-α and IFN-β after the virus challenge. These findings are following a recent study that demonstrated the absence of IFN induction in the alveolar macrophages challenged with SARS-CoV-2 (Dalskov *et al*., 2020). A possible explanation is the presence of a cap structure on the viral genome. The *Coronoviridae* family contains this structure enabling the virus to evade recognition by PRRs and prevent the host innate immune response mediated by the RIG-I/mitochondrial antiviral-signaling (MAVS) pathway that recognizes single-stranded RNAs without a cap structure (Chen & Guo, 2016; Pichlmair *et al*, 2006). Moreover, different SARS-CoV-2 proteins were found to antagonize type I IFN production through other mechanisms (Xia *et al*, 2020). SARS-CoV-2 nonstructural protein 6 (nsp6) binds TANK binding kinase 1 (TBK1) to suppress IRF3 phosphorylation, and nsp13 binds and blocks TBK1 phosphorylation. In addition, open reading frame 6 (ORF6) binds importin Karyopherin α 2 to inhibit IRF3 nuclear translocation, and ORF7b prevents STAT1 phosphorylation and nuclear translocation, consequently inhibiting the transcription of multiple IFN stimulated genes (ISGs) which possess antiviral functions (Xia *et al*., 2020). In parallel, the negative fold change of IFN-related interaction networks: “IFN signaling and positive regulation of RIG-I signaling pathway” and “IFN-α/β signaling and IFN-γ signaling” indicates dysregulated IFN responses in the SARS-CoV-2-challenged MDMs. Thus far, our data suggest an aberrant IFN response induced by SARS-CoV-2 in human MDMs. Early IFN induction was halted at the transcriptional level one day post-infection and did not proceed to the translational level. Thus, we postulate that aberrant IFN responses in the face of a robust inflammatory environment presage a lack of viral infection control and multiorgan damage in COVID-19 (Gavriatopoulou *et al*., 2020).

In summary, we demonstrate that abortive SARS-CoV-2 macrophage infection triggers a unique signature of inflammatory responses. This includes combinations of abortive infection with delayed viral clearance, excessive pro-inflammatory cytokine and chemokine productions, and attenuated IFN responses. Each and all imply dysfunction of the innate immune system after viral exposure. Transcriptomic and proteomic profiles of SARS-CoV-2-challenged MPs provide signatures for understanding COVID-19 pathogenesis. The lack of adequate antiviral innate immune responses in the midst of cytokine storm heralds an absence of viral infection control that exacerbates clinical manifestations and contributes to end-organ damage for COVID-19-related morbidities and mortalities.

## Data availability

This study includes no data deposited in external repositories.

## Acknowledgments

The authors thank the UNMC Elutriation and Cell Separation Core Facility (Myhanh Che and Na Ly) for providing human monocytes. The authors would also like to thank Dr. Pawel Ciborowski and his research group within the Department of Pharmacology and Experimental Neuroscience at UNMC for discussion and assistance with the proteomic sample preparation. The authors acknowledge the UNMC Mass Spectrometry and Proteomics Core Facility for assistance with the proteomic analysis. The authors would also like to acknowledge the UNMC Flow Cytometry Research Core Facility for assistance with the flow cytometric analysis. The authors would like to thank Dirk Anderson (The Biotech Microscopy Core Research Facility of University of Nebraska-Lincoln) for his assistance with TEM sample preparations.

## Author contributions

M.M.A.: Investigation, Methodology, Validation, Visualization, Formal analysis, Data curation, Writing-original draft, review and editing; P.Y.: Investigation, Methodology, Formal analysis, Data curation, Writing-review and editing; J.M.: Investigation, Methodology, Data curation, Writing-review and editing; K.E.O.: Validation, Visualization, Writing-review and editing; F.S.: Investigation, Validation, Writing-review and editing; Y.Z.: Investigation, Validation, Data curation, Writing-review and editing; L.J.: Investigation, Formal analysis, Writing-review and editing; K.P.: Investigation, Validation, Writing-review and editing; A.A.: Investigation, Validation, Visualization, Writing-review and editing; S.N.B.: Supervision, Visualization, Writing-review and editing; R.L.M.: Conceptualization, Methodology, Supervision, Writing-review and editing; H.E.G.: Conceptualization, Funding Acquisition, Methodology, Resources, Supervision, Writing-original draft, review and editing; All authors have read and approved the final version of the manuscript.

---

**Supplementary Figure 1.**
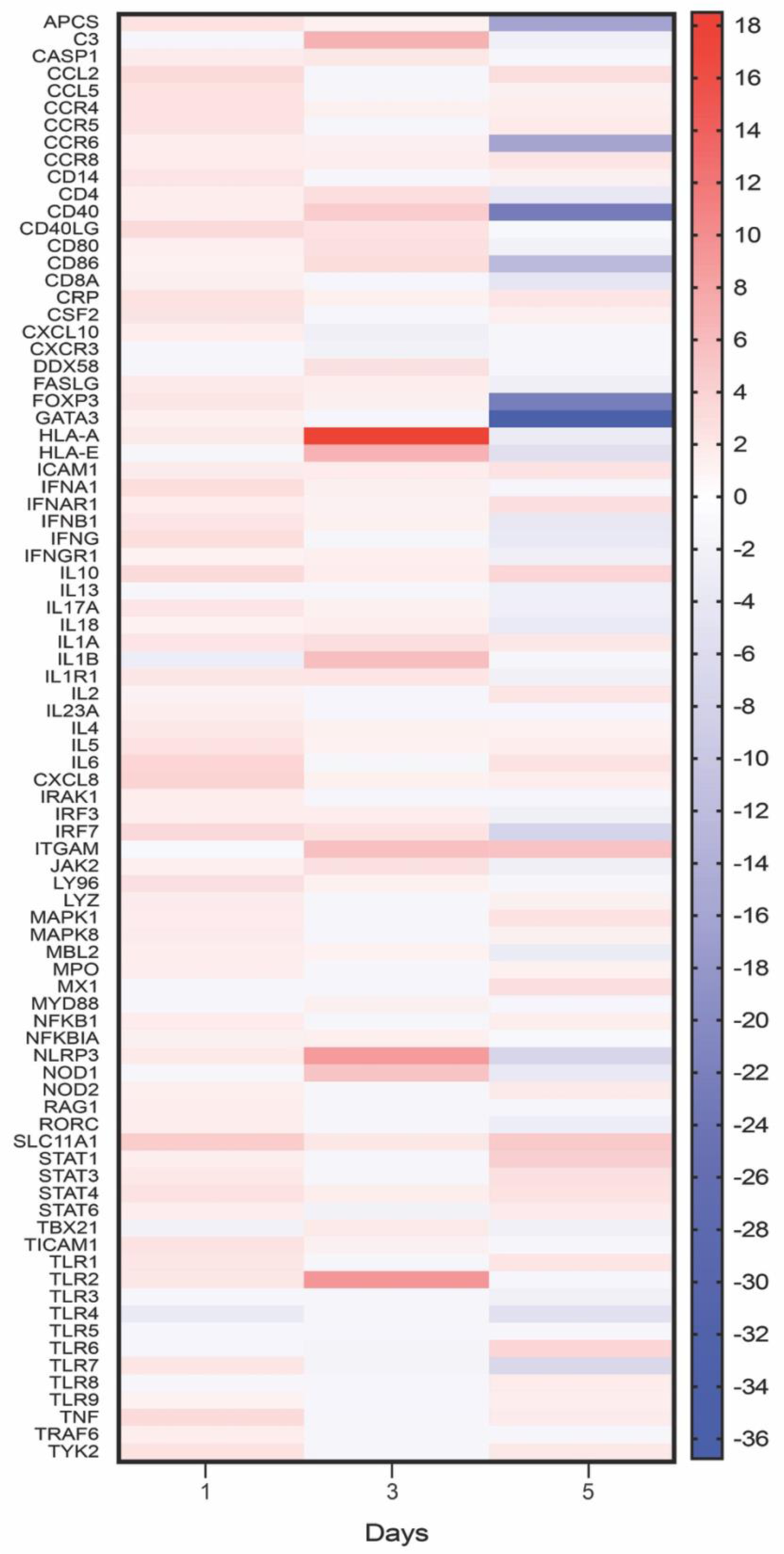
Fold changes of immune response genes in SARS-CoV-2-challenged MDMs. Heat map of fold changes in the expression of 84 genes specific for human innate and adaptive immune responses in SARS-CoV-2-challenged MDMs compared to mock-challenged MDMs at different time points after the infection, determined using RT2 Profiler Human Innate and Adaptive Immune Response 96-well Array. Fold changes were determined via Qiagen’s RT2 Profiler analysis software (n=4).

**Supplementary Figure 2.**
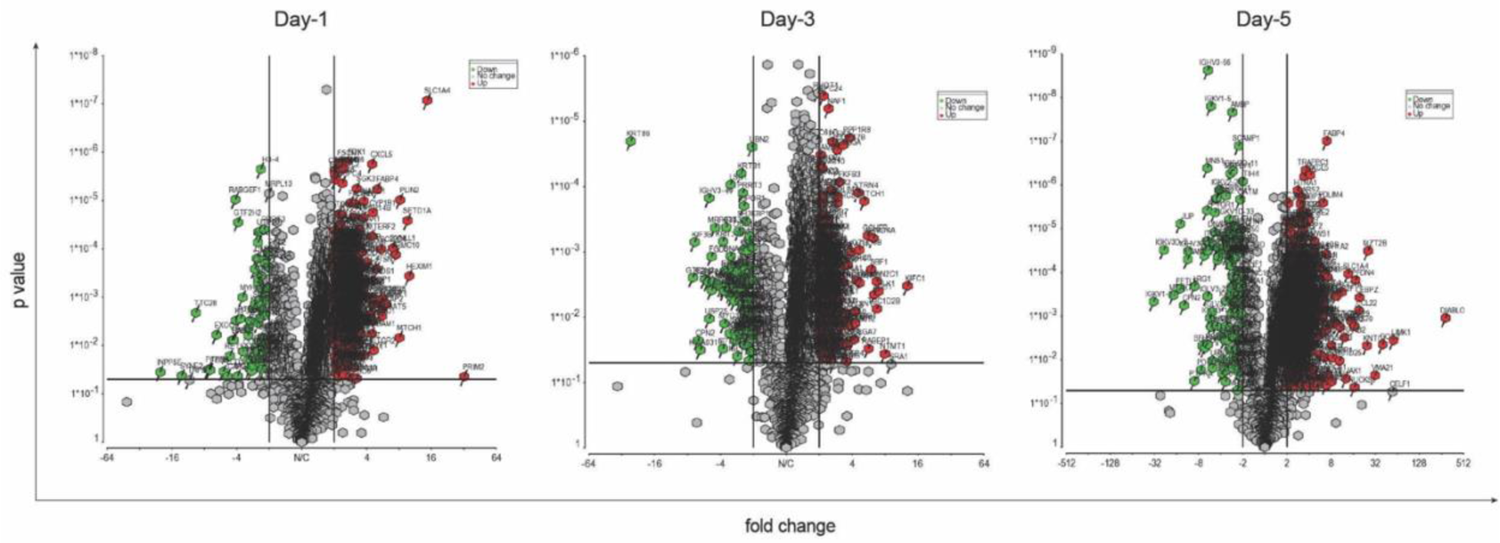
Differential proteomic analysis of SARS-CoV-2-challenged MDMs. Volcano plots showing the fold change plotted against the P value highlighting significantly changed proteins (red – upregulation and green – downregulation; p ≤ 0.05 and an absolute fold change ≥ 2) in SARS-CoV-2-challenged MDMs compared to mock-challenged MDMs at different time points (n=4). The vertical lines correspond to the absolute fold change of 2, and the horizontal line represents a p value of 0.05.

